# PrgE: an OB-fold protein from plasmid pCF10 with striking differences to prototypical bacterial SSBs

**DOI:** 10.1101/2024.03.13.584862

**Authors:** Annika Breidenstein, Anaïs Lamy, Cyrielle P. J. Bader, Wei-Sheng Sun, Paulina H. Wanrooij, Ronnie P-A Berntsson

**Author notes:** Correspondence should be addressed to R.P-A.B.

## Abstract

A major pathway for horizontal gene transfer is the transmission of DNA from donor to recipient cells via plasmid-encoded Type 4 Secretion Systems (T4SS). Many conjugative plasmids encode for a single-stranded DNA-binding protein (SSB) together with their T4SS. Some of these SSBs have been suggested to aid in establishing the plasmid in the recipient cell, but for many their function remains unclear. Here, we characterize PrgE, a proposed SSB from *Enterococcus faecalis* plasmid pCF10. We show that PrgE is not essential for conjugation. Structurally, it has the characteristic OB-fold of SSBs, but it has very uncharacteristic DNA-binding properties. Our DNA-bound structure shows that PrgE binds ssDNA like beads on a string, and this plasticity of PrgEs oligomerization is further confirmed by *in vitro* studies. Unlike other SSBs, PrgE binds both double- and single-stranded DNA equally well. This shows that PrgE has a quaternary assembly and DNA-binding properties that are very different from the prototypical bacterial SSB, but also different from the eukaryotic SSBs.

## Introduction

Horizontal gene transfer is an important way for bacteria to spread genetic information between populations, for example for the propagation of antibiotic resistance or virulence genes^1^. Conjugation is one type of horizontal gene transfer which allows for the transfer of plasmids from donor to recipient cells via Type IV Secretion Systems (T4SS)^2^. These systems are increasingly well-understood in Gram-negative bacteria, where recent cryo-EM structures provide an understanding of the mating channel at a molecular level^3,4^. In contrast, our current understanding of Gram-positive T4SSs is much more limited as such detailed information is not available^5^.

One of the best studied Gram-positive T4SS is from the conjugative plasmid pCF10^6,7^. This plasmid is a clinical isolate from *Enterococcus faecalis*, a commensal pathogen that often causes hospital-acquired infections and is frequently multiresistant to antibiotics^8–11^. pCF10 is a pheromone inducible plasmid with a complex regulation^12,13^. All T4SS proteins on pCF10 are encoded on a single operon, controlled by the P_Q_ promotor. This operon thus contains the genes that code for i) some of the regulatory proteins, ii) the adhesin proteins that facilitate mating pair formation, iii) the proteins that form the mating channel and iv) the DNA-transfer and replication (Dtr) proteins, including ATPases that provide the energy for DNA transport, and the relaxosome proteins PcfF and PcfG, which aid in processing the plasmid DNA prior to transfer (**Fig. 1**)^5,14^.

**Figure 1.**
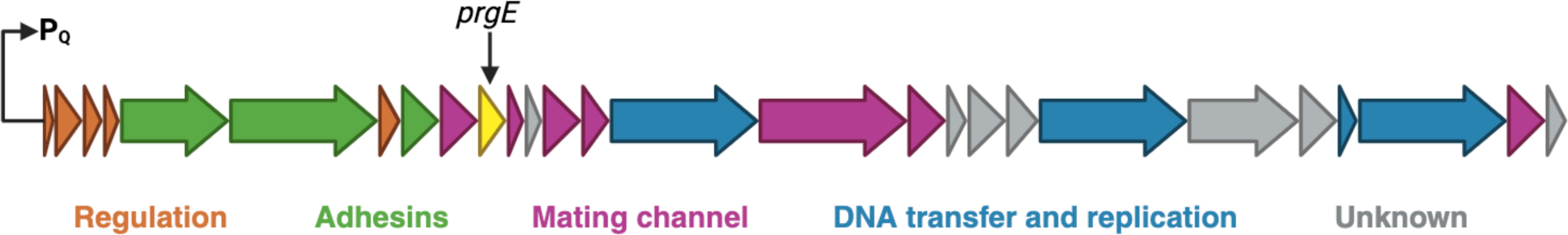
Schematic overview of the genes included in the P_Q_ operon of pCF10. Each arrow represents one gene, coloured by its proposed function in the T4SS. Genes coding for proteins involved in T4SS regulation are shown in orange, surface adhesins in green, mating channel in purple, DNA transfer and replication (Dtr) proteins in blue, and genes of unknown function in gray. The length of the arrows is approximately to scale of the corresponding genes. *prgE* is highlighted in yellow.

Many conjugative plasmids encode additional proteins that are not directly involved in conjugation, but have various functions that confer competitive advantages to the plasmid^15^. PrgE is a small soluble protein that is encoded roughly one third into the P_Q_ operon, in between genes encoding for the mating channel (**Fig. 1**). PrgE has not been previously characterized and its role in type IV secretion is therefore unknown, but it has been suggested that PrgE is a single-stranded DNA-binding protein (SSB), based on its sequence homology of 37 % to a SSB in a lactococcal phage^6,16^.

SSBs are involved in all molecular mechanisms that require manipulation of single-stranded (ss) DNA, such as DNA-replication, recombination, and repair and can be found in all kingdoms of life^17^. Generally, SSBs share a structural motif, the oligosaccharide/oligonucleotide-binding (OB)-fold. The motif consists of a 5-stranded beta-barrel followed by a single alpha-helix. However, there is a lot of variability in the loops between the beta-strands and the length of OB domains can range from 70 to 150 amino acids and they often have a low primary sequence identity of 5-25 %^18,19^. While the topology of the OB-fold is well conserved, the quaternary organization of SSBs varies between the different kingdoms of life. The *E. coli* SSB, which is the prototype for bacterial SSBs, forms a homotetramer with two distinct DNA-binding modes, depending on salt and protein concentrations. In the first binding mode, *E. coli* SSB interacts with ssDNA with only two of its subunits, while the ssDNA wraps around the full tetramer in the second^20–22^. In eukaryotes, the prototypical SSB is replication protein A (RPA). RPA forms a heterotrimer consisting of RPA70, RPA32 and RPA14, with corresponding molecular weights, with each subunit containing at least one OB-fold^23,24^. When it comes to archaea, some phyla have SSBs that resemble bacterial SSBs, while others have more in common with eukaryotic RPA^25^. Some viruses rely exclusively on host SSBs, while others encode their own proteins, with a large diversity of characteristics, some of which act as monomers^26,27^. However, there is also variation within the kingdoms, as many bacterial and eukaryotic species have more than one type of OB-fold protein, which can vary significantly from their respective prototypes^27–30^.

In addition to chromosomal SSBs, many prokaryotes also carry conjugative plasmids that encode SSBs^31,32^. These are believed to contribute to plasmid maintenance, and are thought to be important for protecting ssDNA during conjugation^31,33,34^. Many plasmid SSBs can complement for deficiencies in genomic SSB^32^. Recently, it was shown that F plasmid encoded T4SS can translocate plasmid SSB into recipient cells where they function to suppress the mating-induced SOS-response^15,35^. However, it is not known whether SSBs encoded on conjugative plasmids from Gram-positives are functionally analogous.

In this study, we show that PrgE plays no essential role in conjugation, but that it has very unusual DNA-binding properties. Crystal structures of apo and DNA-bound PrgE show that PrgE has the characteristic OB-fold of SSBs, but that it binds ssDNA in a filamentous way, which is further supported by *in vitro* experiments. We also present data that shows that PrgE unexpectedly binds both ssDNA and dsDNA equally well.

## Materials and methods

### Cloning, plasmids and strains

Strains, oligos and plasmids used in this study are listed in **Table S1**. *E. coli* strains were cultured in Lysogeny Broth (LB) or Terrific Broth (TB) supplemented, when necessary, with antibiotics at the following concentrations: 100 µg/mL kanamycin, 20 µg/mL gentamycin and 25 g/mL chloramphenicol. *E. faecalis* strains were cultured in Brain-Heart-Infusion (BHI) broth or Tryptic Soy Broth without Dextrose (TSB-D) supplemented, when necessary, with antibiotics at the following concentrations, 10 µg/mL chloramphenicol, 10 µg/mL tetracycline, 25 µg/mL fusidic acid, 20 µg/mL erythromycin.

The sequence encoding *prgE* was PCR-amplified from the pCF10 plasmid using primers PrgE_FX_F and PrgE_FX_R and cloned into the intermediate vector pINIT_kan after digestion by *SapI*, using the FX cloning system^36^. It was sub-cloned into the expression vector p7XC3H, which provides a C-terminal 10xHis-tag and a 3C protease cleavage site, before transformation of *E. coli* ArcticExpress (DE3) cells. The sequence encoding *pcfG* was PCR-amplified using the primers PcfG_F and PcfG_R and cloned into a pET24d vector after digestion with *Eco31I*, which provides a N-terminal 10xHis-tag and a SUMO-tag, before transformation into *E. coli* BL21 (DE3) cells.

The *E. faecalis* PrgE deleted strain, OG1RF:pCF10Δ*prgE*, was obtained by allelic exchange and counter-selection using a pCJK218 plasmid^37^, leaving the nucleotides encoding the first and last five amino acids of the protein. About 800 bp of the upstream and downstream region of PrgE were PCR-amplified using the primer pairs PrgE-UF-F/PrgE-UF-R and PrgE-DF-F/PrgE-DF-R, respectively. The products were digested by *BamHI/SalI* for the upstream region and SalI/NcoI for the downstream region, prior to cloning into the pCJK218 digested by *BamHI/NcoI*. The resulting plasmid was used to transform *E. faecalis* OG1RF:pCF10 by electroporation^38^. The PrgE deleted transformants were obtained by switching temperature to induce allelic exchange as described by Vesić and Kristich^37^, and the gene deletion was subsequently confirmed by sequencing.

### Protein production

Proteins were expressed using the LEX system (Large-scale EXpression system, Epiphyte 3). PrgE was transformed in *E. coli* ArcticExpress (DE3) cells and cultivated in TB medium supplemented with 0.4 % glycerol. The cultures were grown at 30°C until an OD_600_ of 0.8, then, cooled down to 12°C before 0.4 mM IPTG was added to induce protein expression. After 24 h, cells were centrifuged at 4000 x*g* during 20 min. PcfF was produced the same way, with the exception that BL21 (DE3) cells were used, and cultures were grown at 37°C before lowering the temperature to 18°C prior to induction, and harvested after 20 h. PcfG was produced in Origami (DE3) cells using autoinduction TB media. Cultures were grown at 37°C until OD 0.6 was reached, followed by 24 h at 25°C without the addition of IPTG.

### Protein purification

Cell pellets were resuspended in different lysis buffers. For PrgE this lysis buffer consisted of 50 mM Tris-HCl pH 8, 10 % glycerol, 500 mM NaCl, 10 mM imidazole, 0.2 mM AEBSF, 1 mM DTT, 1 mM MgSO_4_ and 0.02 mg/mL DNase I. For PcfF the lysis buffer was 50 mM HEPES pH 7.5, 500 mM NaCl, 0.2 mM AEBSF and 0.02 mg/mL DNase I. For PcfG the lysis buffer was 50 mM Tris-HCL pH 8, 10 % glycerol, 500 mM NaCl, 10 mM imidazole, 0.2 mM AEBSF and 0.02 mg/mL DNase I. Resuspended cells were lysed in a Cell Disruptor (Constant Systems) at 25 kPsi and centrifuged at 30,000 x *g* for 30 min at 4°C.

PrgE-His supernatant was incubated for 1 hour at 4°C with gentle rocking with Ni-NTA resin (Protino®). After incubation, the mix was transferred into a gravity flow column and washed with three subsequent 20 CV washes with wash buffer (20 mM Tris-HCl pH 7.5, 5 % glycerol, 500 mM NaCl, 50 mM imidazole), LiCl-wash buffer (wash buffer supplemented with 2 M LiCl) and wash buffer. The resin was then incubated overnight at 4°C with gentle rocking in elution buffer (20 mM Tris-HCl pH 7.5, 500 mM NaCl, 5 % glycerol, 50 mM imidazole, 1 mg of PreScission protease, 1 mM DTT), during which the His-tag was cleaved. The flow through, as well as an additional wash with 5 CV elution buffer, was collected and concentrated using Amicon Ultra Centrifugal filters with a molecular weight cutoff of 10 kDa. The protein was then subjected to size exclusion chromatography (SEC) in 20 mM HEPES pH 7.5 and 300 mM NaCl on a HiLoad 16/600 Superdex 200 pg column using an Äkta Purifier (Cytiva). For buffer exchange before various experiments, PrgE was subjected to a second SEC using a Superose 6 Increase 10/300 GL column on an Äkta Pure (Cytiva).

GST-PcfF supernatant was incubated for 1 h with Glutathione resin (GE Healthcare) at 4°C and subsequently washed with 50 CV wash buffer (20 mM HEPES pH 7.5, 200 mM NaCl) prior to elution with 20 mM HEPES pH 7.5, 200 mM NaCl, 30 mM glutathione. The protein was concentrated with Amicon Ultra Centrifugal filters with a molecular weight cutoff of 10 kDa prior to SEC in 20 mM HEPES pH 7.5, 200 mM NaCl on a Superdex 200 Increase 10/300 GL column using an Äkta Pure (Cytiva).

His-PcfG supernatant was incubated for 1 h at 4°C with pre-equilibrated HisPur^TM^ Cobalt Resin (Thermo Scientific) to three subsequent 20 CV washes with wash buffer (20 mM HEPES pH 7.5 5 % glycerol, 300 mM NaCl, 30 mM imidazole), LiCl-wash buffer (wash buffer supplemented with 2M LiCl) and wash buffer. PcfG was then eluted in 15 CV elution buffer (20 mM HEPES pH 7.5 5 % glycerol, 300 mM NaCl, 150 mM imidazole). The protein was loaded onto a HiTraP Heparin HP (5 mL) column (GE Healthcare) equilibrated with Buffer A (20 mM HEPES pH 7.5, 150 mM NaCl). PcfG was eluted in a salt gradient to 100 % Buffer B (20 mM HEPES pH 7.5, 1000 mM NaCl).

### Crystallization and structure determination

SEC purified PrgE, with a concentration of 11 mg/mL, was used for crystallization trials. Crystals appeared after 2-5 days, at 20°C, using the vapor diffusion method in a condition with 0.2 M LiSO_4_, 0.1 M K Phos Cit pH 4.2, 20 % w/v PEG 1000 in a 2:1 ratio. For the DNA-bound structure, 117 µM of single-stranded poly-A 60mer was added to 6 mg/mL PrgE and mixed in a 1:2 ratio with a reservoir solution containing 15 % v/v PEG 400, 50 mM MES pH 6.5, 80 mM Mg acetate, 15 mM MgCl_2_. Crystals were flash-frozen in liquid nitrogen without additional cryo-protectant. X-ray diffraction data were collected at the ID30A-3 (apo) or ID23-1 (DNA-bound) beamlines at the ESRF, France and processed using XDS^39^. The space group of both crystals was P2_1_2_1_2_1_ and the phase problem was solved in Phenix Phaser^40^ using molecular replacement with an AlphaFold2^41^ model of PrgE where the flexible extremities of the protein had been removed, generated using ColabFold version 1.5.2 using default settings^42^. The asymmetric unit of the crystal contained two copies of PrgE for the apo structure. The asymmetric unit of the DNA-bound protein contained three copies of the protein and a 15 nucleotide stretch of the single-stranded DNA. The chosen asymmetric unit thus contains only a quarter of the full ssDNA that the protein was crystallized with. We chose to do so since the ssDNA has continuous density throughout the crystal packing, and this greatly simplified the refinement process. The structures were built in Coot^43^ and refined at 2.7 Å using Refmac5^44^ and obtained R_work_/R_free_ values of 23.45 and 27.77 for the apo structure and 23.05 and 25.23 for the DNA-bound structure. Further refinement statistics can be found in **Table S2**. Atomic coordinates and structure factors have been deposited with the Protein Data Bank with the accession codes 8S4S and 8S4T for the apo and DNA-bound structures, respectively.

### SEC-MALS

For analysis of the oligomeric state of PrgE, 150-300 µL of 1 mg/mL PrgE (with a theoretical mass of 17 kDa) was loaded on a Superdex 200 Increase 10/300 GL column, equilibrated in buffer (20 mM HEPES pH 7.5 and 300 mM NaCl) via an ÄKTA Pure (Cytiva) that was coupled to a light scattering (Wyatt Treas II) and refractive index (Wyatt Optilab T-Rex) detector to determine the molecular weight of the elution peak via multi-angle laser light scattering (SEC-MALS). Data was analyzed using Astra software (version 7.2.2; Wyatt Technology).

### Crosslinking

PrgE crosslinking experiments were performed by incubating 30 µg of protein with 2 mg of disuccinimidyl suberate (DSS) in 20 mM HEPES pH 7.5 and 300 mM NaCl for 30 min at 20°C. The reaction was quenched by adding 100 mM Tris/HCl pH 8.0 at least 10 min prior to analysis using SDS-PAGE with Coomassie Brilliant Blue staining.

### Preparation of DNA substrates

Oligonucleotides were purchased from Eurofins and are listed in **Table S1**. For double-stranded substrates, one nmol of each oligonucleotide was annealed to an equimolar amount of its complementary strand by denaturing at 95°C for 5 min in TE buffer (50 mM Tris-HCl pH 8.0, 1 mM EDTA) containing 100 mM NaCl, and allowing the reaction mixture to cool to room temperature. The DNA was separated on a 15 % acrylamide gel in 0.5 × TBE (15 mM Tris, 44.5 mM boric acid, 1 mM EDTA), stained with 3 × GelRed (Biotium) for 30 min and visualized by using Chemidoc^TM^ (Bio-Rad). The bands corresponding to double-stranded molecules were excised with a clean razor blade, eluted from crushed gel slices into TE buffer (10 mM Tris-HCl, pH 8.0, 1 mM EDTA) and purified by phenol-chloroform extraction and isopropanol precipitation.

### Fluorescence anisotropy assay

Single-stranded and double-stranded oligonucleotides of 30 nt or 60 nt with a 5’ FITC label were diluted to 20 nM in binding buffer (20 mM HEPES pH 7.5, 50 or 100 mM NaCl, as indicated). Before use, the single-stranded oligonucleotides only were boiled for 5 min at 95°C and chilled on ice. Fluorescence anisotropy reactions containing 10 nM oligonucleotide and 0 - 20 *μ*M PrgE in binding buffer were pipetted in duplicates onto black shallow 384-well microplates (OptiPlate-F, PerkinElmer) and incubated in the dark for 30 min at room temperature. Fluorescence intensities were collected from above on a CLARIOstar® *Plus* plate reader (BMG Labtech) with the excitation and emission wavelengths 480 nm and 520 nm, respectively. Fluorescence anisotropy in millianisotropy units (mA) was calculated using MARS Data analysis Software (BMG Labtech) according to Equation 1:

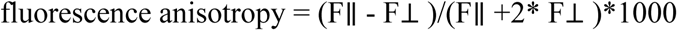

where F∥ and F⊥ are the parallel and perpendicular emission intensity measurements corrected for background (buffer). PrgE alone exhibited no fluorescence. The dissociation constant (K_d_) was determined by fitting data to a quadratic equation by non-linear regression analysis in GraphPad Prism software (GraphPad Software, Inc., USA) using Equation 2:

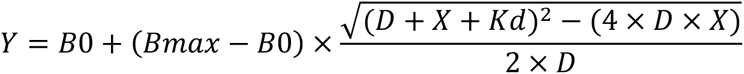

where Y is the anisotropy value at protein concentration X, X is the concentration of PrgE in *μ*M, B0 and Bmax are specific anisotropy values associated with free DNA and total DNA-PrgE respectively, and D is the concentration of DNA in *μ*M.

### Pull-down experiments with relaxosome components

PrgE pull-down experiments were performed in 20 mM HEPES pH 7.5 and 200 mM NaCl by mixing either 2 nmol GST-PcfF or PcfG-His (baits) with 4 nmol PrgE without tag (pray) and 100 µl of the resin (Glutathione resin (GE Healthcare) when using PcfF and Ni-NTA (Protino®) for PcfG). The proteins were incubated for 15 min at 4°C prior to collecting the flow through and washing with 5 x 5 CV wash buffer and eluting with 2 x 5 CV elution buffer. For GST-PcfF pull-downs, 20 mM HEPES pH 7.5 and 200 mM NaCl was used as wash buffer and 20 mM HEPES pH 7.5, 200 mM NaCl and 30 mM Glutathione as elution buffer. For His-PcfG pull-downs, wash buffer contained 20 mM HEPES pH 7.5, 200 mM NaCl, 30 mM imidazole and elution buffer 20 mM HEPES pH 7.5, 200 mM NaCl, 500 mM imidazole. The samples were analyzed on SDS-PAGE and stained with Coomassie Brilliant Blue.

### Conjugation assays

Donor (OG1RF:pCF10 or OG1RF:pCF10Δ*prgE*) and recipient (OG1ES) strains were inoculated with the indicated antibiotics and incubated overnight at 37°C with agitation. The next day, the overnight cultures were refreshed in BHI media without antibiotics in a 1:10 ratio. For conjugation assays in exponential phase, cells were directly induced to express the T4SS with 5 ng/mL cCF10 for 1 h at 37°C without agitation. For conjugation assays in stationary phase, cultures were first incubated for 3 h at 37°C with agitation prior to induction. Donor and recipient cells were then gently mixed in a 1:10 ratio and incubated for 30 min at 37°C without agitation. To disrupt the ongoing conjugation, cells were vortexed and placed on ice for 10 min.

A serial dilution was performed with cold media and 10 µl of the appropriate dilutions were spotted in triplicates on the top of a square BHI agar plate and placed in an upright position to allow the drops to run down the plate to facilitate counting of the colonies. To select donor cells, BHI agar contained 10 µg/mL tetracycline and 25 µg/mL fusidic acid and to select for transconjugant cells, BHI agar contained 10 µg/mL tetracycline and 20 µg/mL erythromycin. The plates were incubated for approximately 24 hours at 37°C before colonies were counted and enumerated for colony forming units (CFU). Conjugation efficiency was determined as CFU of transconjugant over CFU of donor (Tc’s/Donors). Experiments were done in triplicates and are reported with their standard deviation.

For the serial passaging, conjugation assays were performed in exponential phase as described above. Three colonies of the transconjugant plates from passage 1 were picked to start new overnight cultures, that were then used as donor cells for the following passage. In passage 2, donor cells were therefore OG1ES:pCF10, and OG1RF without a plasmid served as recipient cells. Three trans-conjugant colonies from passage 2 served as donor cells for passage 3 with OG1ES as recipient cells and trans-conjugants cells from passage 3 were donors for passage 4 with OG1RF as recipient. Donor and transconjugant cells were selected as previously described for passage 1 and 3. For passage 2 and 4, BHI agar containing 10 µg/mL tetracycline and 20 µg/mL erythromycin was used to select for donor cells and BHI agar containing 10 µg/mL tetracycline and 25 µg/mL fusidic acid was used to select for transconjugants.

All *in vivo* data is from three biological replicates and was plotted with their standard deviation using GraphPad Prism (version 10.2) (GraphPad Software). Statistical significance was analyzed with One-way Anova.

## Results

### PrgE is not a homolog of genome-encoded E. faecalis SSB

We investigated how similar PrgE is to genome-encoded *E. faecalis* SSB by creating AlphaFold2 models of both proteins. Genomic SSB strongly resembles typical bacterial SSBs and the model aligns with *E. coli* SSB with an RMSD of 0.59 Å over 83 residues (**Fig. S1A**). In contrast, the PrgE model differs significantly. It superimposes with an RMSD of 5.4 Å over 80 residues to the model of genome-encoded *E. faecalis* SSB, with differences in the part of the beta-sheet that is involved in DNA-binding in typical bacterial SSBs (**Fig. S1B**). To compare PrgE to other proteins, we performed sequence-based homology searches, which yielded very little insight, besides that PrgE is thought to be an SSB. Instead we performed a structural homology search using Foldseek^45^. However, also using this methodology the top hits were only distantly related proteins with an OB-fold, with high E-values or low TM scores (**Table S3**). This indicates that the structure of PrgE is different than that of previously studied SSBs.

### PrgE has an OB-fold

PrgE was produced in *E. coli* and purified to homogeneity. We solved the crystal structure of apo PrgE to 2.7 Å, using the AlphaFold2 model of PrgE as a template for molecular replacement. The asymmetric unit contained two copies of the protein in space group P2_1_2_1_2_1_. Both copies were modeled from residue 1-130, with residues 34 and 35 missing in loop 1 of chain A (**Fig. S2**). For both chains, the remaining C-terminal part (residues 131-144) is missing in the density. PISA analysis shows that this dimer has an interface area of 680 Å^2^, with 9 H-bonds and 3 salt bridges. The overarching fold of the protein corresponds to an oligosaccharide/oligonucleotide-binding (OB)-fold, characterized by 5 beta-strands that form a beta-barrel with a 1-2-3-5-4-1 topology, which is only partially closed between strands 3 and 5 for PrgE (**Fig. 2A**). PrgE also has a 42 residues long region between strands 3 and 4 that forms 2 alpha-helices of which the first seemingly contributes to the opening in the barrel between strands 3 and 5. The apo structure overall aligns very well with the predicted AalphaFold2 model of PrgE, having an RMSD of 0.48 Å over 113 residues.

**Figure 2.**
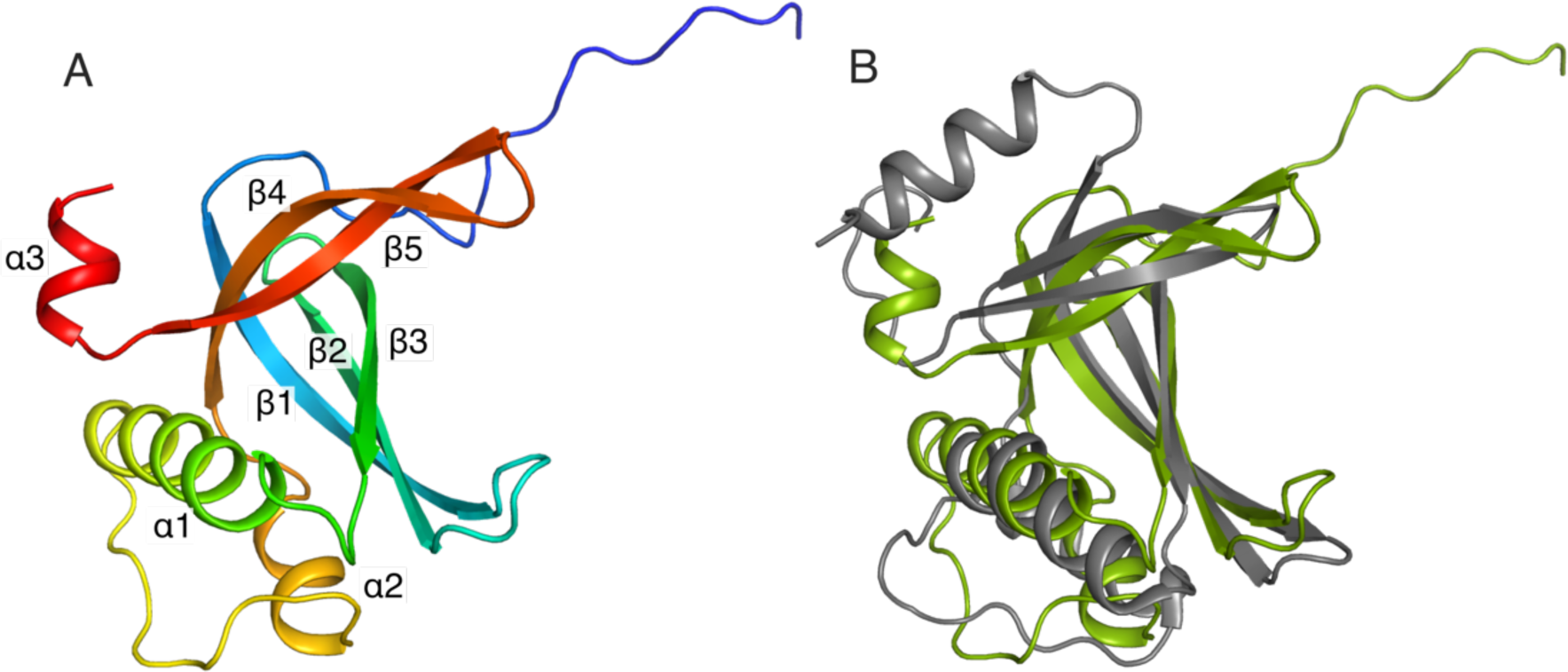
Apo structure of PrgE. **A)** Crystal structure of PrgE colored in rainbow colors from the N-terminus (blue) to the C-terminus (red). All secondary structure elements are marked in the figure. **B)** Superimposition PrgE (green) with the C-terminal domain of RadD (grey, PDB: 7R7J). The beta-sheet superimposes relatively well, but there are larger differences in the orientation of the alpha-helices.

We used DALI^46^ and Foldseek^45^ to search the Protein Data Bank (PDB) for the closest structural homolog to PrgE. As with the previous searches with the AlphaFold2 model, the hits had generally very low scores with E-values in Foldseek being in the 10^−2^ range. The best hit from DALI was the C-terminal domain with unknown function of the *E. coli* helicase RadD^47^ (PDB: 7R7J) with a Z score of 7.1. However, there are substantial structural differences which is highlighted by having an RMSD of 4.02 Å over 104 residues between the two structures (**Fig. 2B**).

### PrgE oligomerizes in vitro

Since the oligomerization of PrgE might be different in solution than in the crystal, we investigated the oligomerization behavior of PrgE *in vitro*. We noticed that the volume at which PrgE eluted on size exclusion chromatography (SEC) differed depending on the salt concentration of the buffer (**Fig. 3A**), as well as the protein concentration (**Fig. 3B**). This indicates that PrgE is able to oligomerize. To gain deeper insight into the oligomeric state, we performed size-exclusion chromatography coupled to multi-angle light scattering (SEC-MALS), with 60 µM PrgE in 300 mM NaCl conditions. The molecular mass of the elution peak was 51.1 +/− 2.8 kDa, which corresponds well to a trimer (the theoretical molecular mass of the PrgE monomer is 17 kDa) (**Fig. 3C**). However, all SEC traces show an asymmetric peak, trailing to the right, indicating the presence of smaller oligomeric species. In addition to this, gentle crosslinking of purified PrgE also captured multiple oligomeric states (**Fig. 3D**). These results show that PrgE can exist in various oligomerization states *in vitro*, and that its oligomerization is both salt-and protein concentration-dependent.

**Figure 3.**
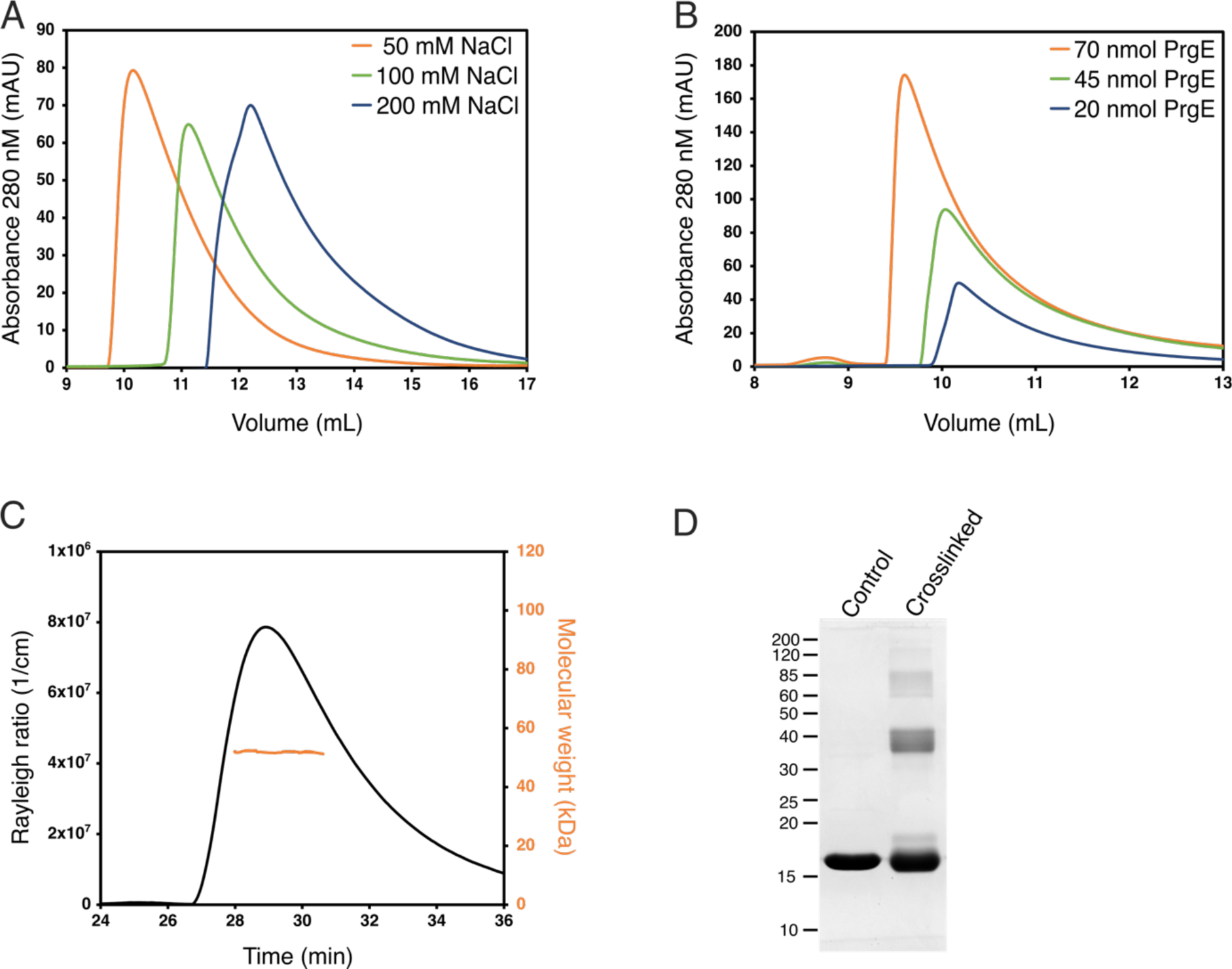
Oligomerization of PrgE. **A)** Size exclusion chromatogram of PrgE (on a Superose 6 column) shows that the elution volume, which is coupled to protein radius, depends on the salt concentration. **B)** Size exclusion chromatogram of PrgE (on a Superdex 200 column), in the same salt concentration but with different protein concentrations, shows that the elution volume decreases with increasing protein concentration. **C)** SEC-MALS analysis of 60 µM PrgE in 300 mM NaCl. The black line, plotted on the left axis, indicates the Rayleigh ratio, which is directly proportional to the intensity of the scattered light in excess of the buffer. The orange line, plotted on the right axis, indicates the molecular weight of the protein measured throughout the peak. The average molecular weight was 51.1 ± 2.8 kDa. **D)** SDS-PAGE of PrgE, with or without crosslinking with DSS.

### PrgE binds ssDNA in a filamentous manner

We also crystallized PrgE together with a single-stranded Poly-A 60-mer DNA in a molar ratio of 1:3. The obtained crystallographic data was refined in space group P2_1_2_1_2_1_ with the asymmetric unit containing three copies of the protein sitting on a string of 15 ssDNA bases. While there are only 15 bases in the asymmetric unit, the ssDNA shows a continuous density throughout the crystal packing (**Fig. S3A**). Compared to the apo structure of PrgE, a few more residues are visible at the C-terminal end (until residues 136 of 144), continuing as an alpha-helix as predicted by the AlphaFold2 model. The DNA does not get wrapped around PrgE, like it does with *E. coli* SSB^20^, rather PrgE interacts with the DNA like beads on a string, with the N-terminal tail of one PrgE binding to the neighboring PrgE, using interactions between polar side chains (**Fig. 4A**). PISA analysis shows that the interaction areas between the PrgE subunits in the DNA-bound structure are between 600-800 Å^2^.

**Figure 4.**
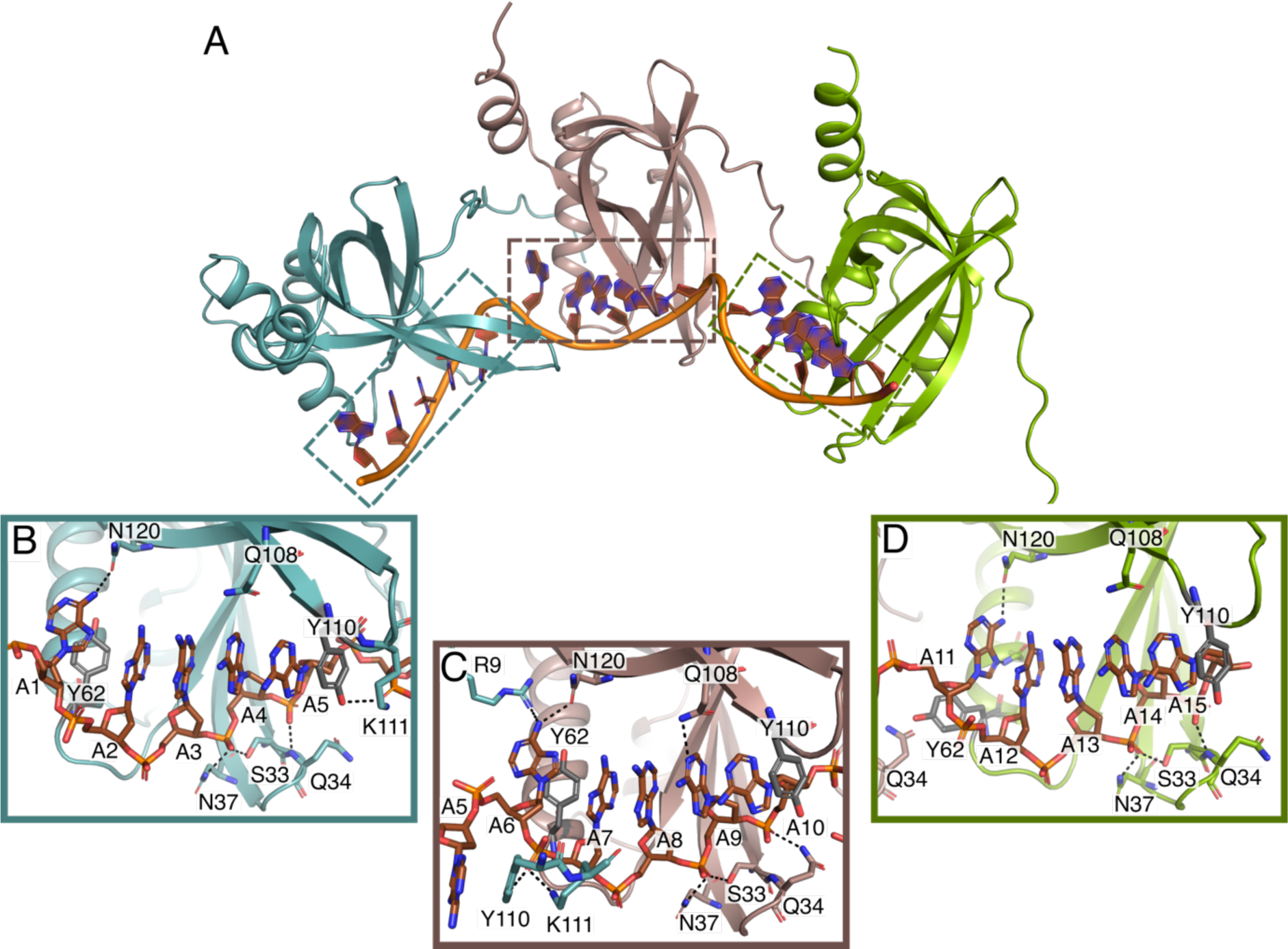
DNA-bound structure of PrgE. **A)** In the asymmetric unit there are three PrgE molecules bound to the ssDNA. **B-D)** Enlarged views of the regions indicated in panel A, highlighting the residues that are important for DNA-binding for each of the three monomers. Black dotted lines show potential hydrogen bonds. The orientation of panels B-D are not the same as in A, to increase clarity and allow easier comparison between B-D.

PrgE binds to the ssDNA between loops 1 and 4, where the beta-barrel is partially open. Each subunit binds to 5 DNA bases. The binding also bends the ssDNA between the protein binding sites, resulting in a kink at every 5^th^ base. The kinks between subunit C’–A and A–B form the same angle. However, the N-terminal tail of chain B bends at a smaller angle and the kink in the DNA chain between subunit B–C is therefore also slightly less pronounced (**Fig. S3B**).

The different PrgE subunits bind to the ssDNA in a similar, but not identical, manner. Many interactions to the phosphate backbone of the ssDNA are the same within all subunits, including with residues Ser33, Gln34 and Asn37 in loop 1 that form H-bonds with the DNA backbone with the 4^th^ and 5^th^ phosphate of each stretch of 5 bases (**Fig 4B-D**). Additional phosphate binding can be found with. Lys111 and Tyr110 in loop 4 in chain A and C, but not B. Interestingly, this loop interacts with the phosphate of the second base of the DNA-binding cassette that is primarily bound by the neighboring copy of PrgE.

In addition to hydrogen-bonding with the phosphate backbone, pi-pi interactions between the aromatic rings of the DNA and two tyrosine residues are of major importance for DNA-binding. Tyr110 stacks on the 5^th^ DNA base in the binding cassette in all subunits. In contrast, the orientation of Tyr62 varies. For chain A and B, Tyr62 points inwards towards the bases, while it is oriented towards the DNA backbone for chain C. Accordingly, the exact orientation of the first DNA base varies between the binding cassettes. In the third binding cassette in the asymmetric unit, base 11 stacks on top of the following 4 bases and forms two H-bonds with PrgE chain C (Asn120 and Asn66). In the other two cassettes (bound to chain A and B) this base is tilted away and only forms one H-bond with Asn120. Other than these interactions with the DNA bases, hydrogen-bonding with DNA bases seems to be less important, consistent with the lack of sequence specificity in DNA-binding. In our structure, only Gln108 of chain B interacts with Adenine 9, with the other copies of Gln108 being close to the DNA but not in hydrogen bonding distance. In conclusion, PrgE binds to ssDNA with a high degree of plasticity.

### PrgE quaternary structure resembles viral SSBs

The overall quaternary structure of PrgE binding to ssDNA is different than that of bacterial or eukaryotic SSBs, where ssDNA commonly wraps around a homotetramer in bacterial SSBs (**Fig. 5A**) and eukaryotic RPA binds DNA as a heterotrimer (**Fig. 5B**). Instead, it appears more similar to that of viral SSBs, which have monomers as a functional unit in DNA-binding (**Fig. 5C**). Each PrgE monomer binds fewer DNA bases (5), which are more neatly stacked on top of each other, compared to other SSBs which have a larger interaction area (**Fig. 5D-F**). The exact DNA-binding mechanisms share some similarities in that stacking interactions with aromatic residues play an important role. However, in PrgE, the responsible residues are tyrosines, while they are phenylalanines and tryptophanes for *E. coli* SSB and RPA, and the viral SSB uses both tyrosines and phenylalanines.

**Figure 5.**
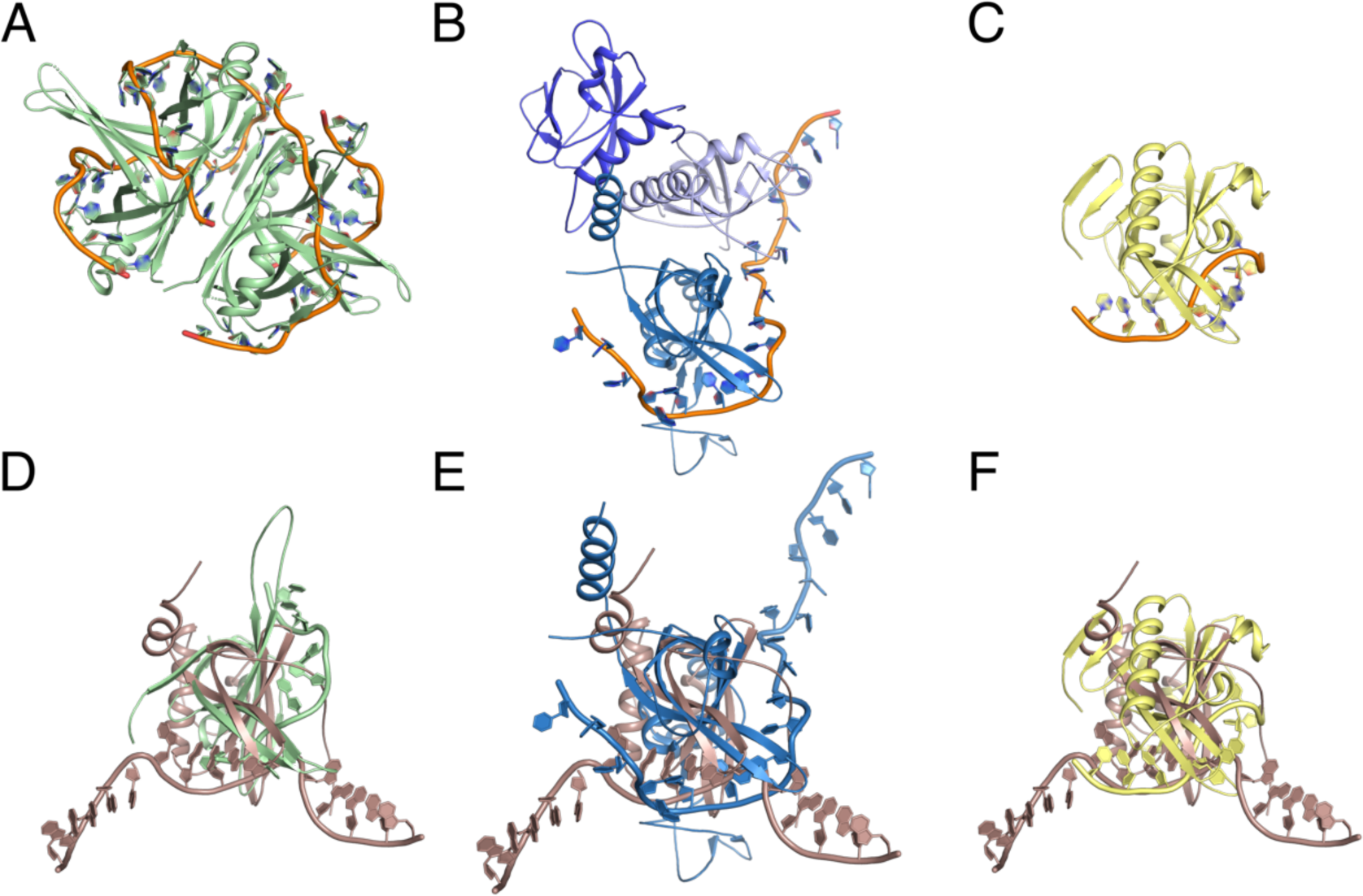
Comparison between PrgE and other SSBs. **A)** *E. coli* homotetrameric SSB bound to ssDNA (PDB: 1EYG). **B)** Yeast heterotrimeric RPA bound to ssDNA (PDB: 6I52). **C)** SSB from Enterobacter phage Enc34 (PDB: 5ODL). **D-F)** Superposition of DNA-bound PrgE with the proteins shown in panels A-C. The view in panel D is rotated 45 degrees on the x-axis when compared to panel A for clarity, the views in panel E-F are the same as in B-C. In panel E, PrgE is aligned to chain C of RPA as it has the highest structural homology to PrgE.

### PrgE binds ssDNA and dsDNA with comparable affinities

Given the suggested function of PrgE as a SSB, we performed binding experiments with the protein on single-stranded (ssDNA) and double-stranded DNA (dsDNA) molecules of 30 or 60 nucleotides. The affinity of PrgE for ssDNA and dsDNA was compared by determining the dissociation constant (K_d_) with each by fluorescence anisotropy (**Fig. 6** & **Fig. S4**). The affinity of PrgE was stronger for the longer 60-mer substrate than for the 30-mer, and higher at lower salt conditions (**Table 1**). Surprisingly, PrgE also bound dsDNA with K_d_ values within the same order of magnitude (K_d_s of 0.33 *μ*M and 0.5 *μ*M for 60-mer ssDNA and dsDNA, respectively, in 50 mM NaCl). These experiments further confirm that the DNA-binding properties of PrgE differ considerably from other SSBs.

**Figure 6.**
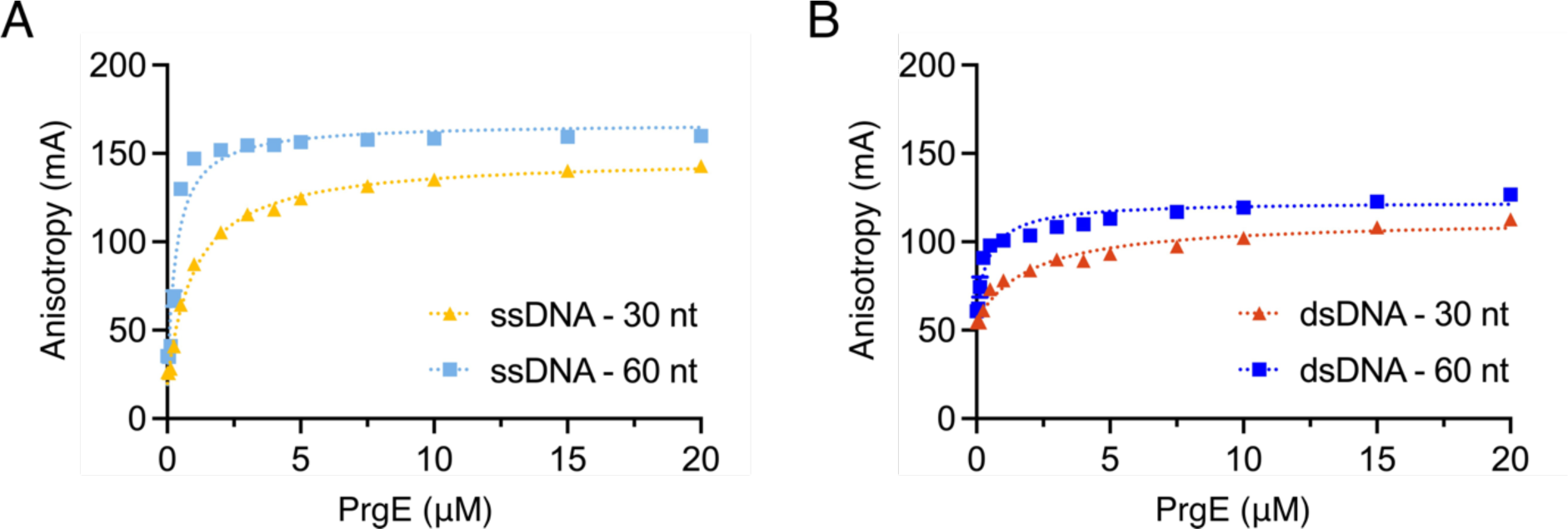
Fluorescence anisotropy measurements of PrgE (0 to 20 *μ*M) in reaction buffer with ssDNA or dsDNA in buffer containing 50 mM NaCl. **A)** PrgE binding to 30-mer (yellow) and 60-mer (light blue) ssDNA substrates. **B)** PrgE binding to 30-mer (orange) and 60-mer (blue) dsDNA substrates. Error bars (only visible over the data point in one instance) represent the standard deviation (n=3).

**Table 1.** The K_d_ values and standard deviations (n=3) for PrgE binding to ssDNA or dsDNA oligonucleotides in 50 mM NaCl as determined by fluorescence anisotropy.

| $K_d \pm SD (\mu M)$ | ssDNA | | dsDNA | |
| --- | --- | --- | --- | --- |
|  | 30 nt | 60 nt | 30 nt | 60 nt |
| 50 mM NaCl | $1.02 \pm 0.02$ | $0.33 \pm 0.02$ | $1.96 \pm 0.26$ | $0.50 \pm 0.14$ |
| 100 mM NaCl | $1.84 \pm 0.15$ | $0.37 \pm 0.01$ | $3.36 \pm 0.28$ | $0.95 \pm 0.03$ |

### PrgE is not essential for conjugation

Given that PrgE is a soluble protein in the T4SS operon that binds DNA, we speculated that it might interact with the DNA transfer and replication proteins PcfF (accessory factor^48^) and/or PcfG (relaxase^49^), which form the relaxosome at the origin of transfer of plasmid pCF10. We therefore conducted pull-down experiments where untagged PrgE was incubated with either the His-tagged PcfG (**Fig. 7A**), or the GST-tagged PcfF (**Fig. 7B**). However, neither of the proteins co-eluted with PrgE, indicating that they do not strongly interact.

**Figure 7.**
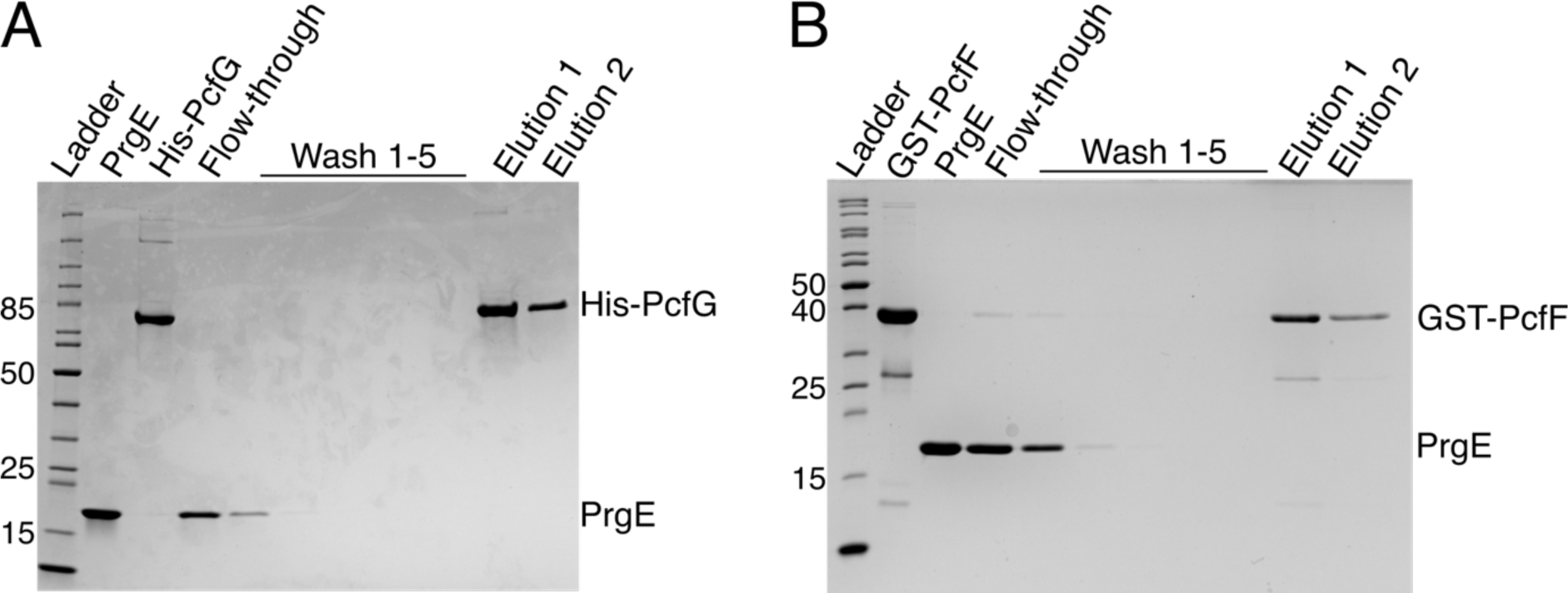
PrgE does not interact with the main components of the pCF10 relaxosome. **A)** Pull-down experiment with the relaxase PcfG, showing the input protein, washes and elution, in which His-PcfG (bait) was unable to pull-down PrgE (pray). **B)** Pull-down experiment in which the relaxosome accessory factor GST-PcfF (bait) was unable to pull-down PrgE (pray).

Since PrgE is likely not part of the relaxosome, we wanted to know if it is essential for conjugation in another way. We therefore created an *E. faecalis* knockout strain (OG1RF:pCF10Δ*prgE*) to explore the function of PrgE *in vivo* by comparing the conjugation efficiency between mutant and wildtype. We tested conjugation both during exponential phase when cells were actively dividing and in stationary phase, where cells are no longer dividing and the availability of other, genome-encoded, SSBs in *E. faecalis* may be different. We observed a decrease in efficiency between exponentially growing cells and cells in stationary phase, but there was no significant difference between Δ*prgE* and wildtype in either condition (**Fig. 8**). We further considered whether multiple conjugative events would be needed to observe an effect. We therefore passaged the plasmids several times between donor and recipient cells, by using trans-conjugant cells as new donor cells. However, also here we did not observe any difference within four passages between Δ*prgE* and wildtype (**Fig. 8**). We conclude that PrgE does not play an essential role in conjugation under the tested conditions.

**Figure 8.**
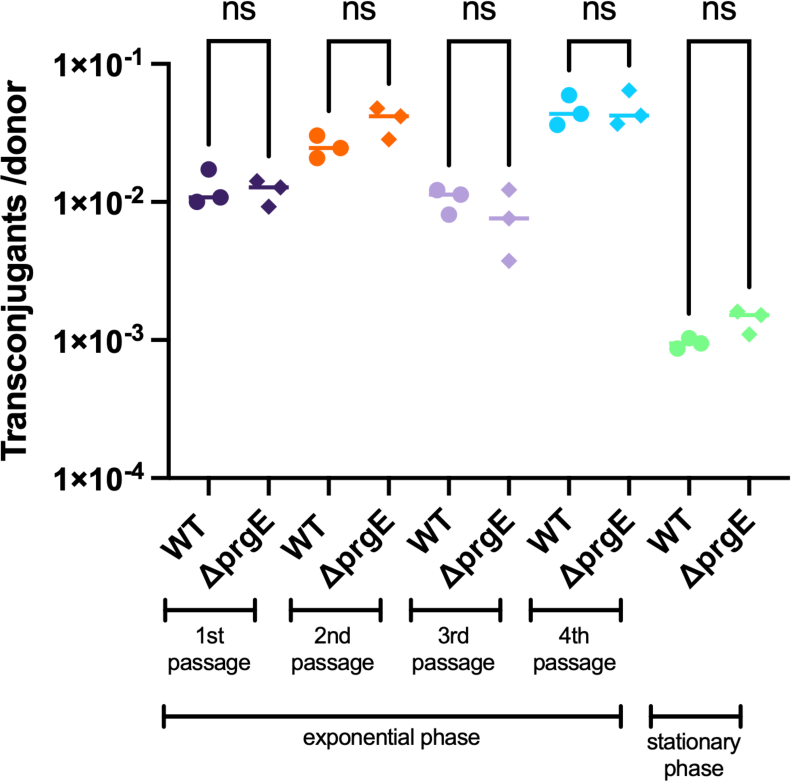
PrgE is not essential for conjugation. Conjugation rates of *E. faecalis* donor cells carrying wildtype pCF10 or pCF10Δ*prgE* either in exponential growth or stationary phase. In exponential growth serial passaging was performed, where transconjugants from one passage were used as donor cells in the following passage. ns stands for not significant.

## Discussion

Many conjugative plasmids, with different incompatibility groups, encode for (at least) one SSB protein, which can often complement for the genome-encoded SSB^32^. In conjugation, SSBs have been proposed to be important for protecting plasmid ssDNA both in donor and recipient cells and to evade the SOS response^33–35,50^. However, all of the available research has been done on SSBs from Gram-negative T4SS system. Here, we characterized the proposed SSB PrgE from the Gram-positive conjugative plasmid pCF10.

By crystallizing PrgE, we showed that it indeed has the typical OB-fold of SSBs, but that its structure has important differenes when compared to other SSB proteins. PrgE has three alpha-helices that are positioned differently from other SSBs, and also differs in its beta-sheet where the DNA-binding regions are. The differences became even more apparent when we analyzed the DNA-bound structure. Each monomer binds DNA in a way that is to be expected, relying on interactions with the DNA backbone and stacking interactions with the bases to achieve DNA-binding in a sequence-independent manner. However, PrgE does not bind DNA as the typical bacterial SSB, which commonly form homotetramers around which they wrap the ssDNA. It is also very different from how eukaryotic SSBs, like RPA, bind ssDNA as heterotrimers. Instead, PrgE binds the ssDNA in a filamentous manner, like beads on a string (**Fig. 4**). Between each binding site the DNA gets bent (**Fig. S3B**). Whether the exact angles are due to crystal packing or are also the ones found in solution is not known. Further supporting the filamentous oligomerization are the different oligomerization states that were observed for PrgE in solution (**Fig. 3**). The oligomerization in the DNA-bound structure is supported by the N-terminal tail of PrgE, which interacts with the neighboring monomer on the DNA-bound structure (**Fig. 4**), a feature that is not found on the prototypical bacterial SSBs.

Our data from the fluorescence anisotropy experiments show a standard hyperbolic binding curve, indicating that the binding is not cooperative in nature (**Fig. 6**, **Fig. S4)**. Surprisingly, we found that PrgE bound dsDNA equally well as ssDNA (**Fig. 6**, **Fig. S4** & **Table 1**). Most characterized SSBs have a high affinity and specificity for ssDNA^27^. As an example, RPA binds mixed ssDNA with affinities of 10-40 nM albeit displaying a preference to pyrimidines, and with K_D_ values to ssDNA up to 3 orders of magnitude lower than to dsDNA^51–53^. To our knowledge, only one studied SSB-like protein shares PrgE’s feature of binding equally well to both ssDNA and dsDNA, namely one from the archaea *Nanoarchaeum equitans*^54^. Given these data, it is clear that PrgE is not a typical SSB, and we therefor refer to it simply as an OB-fold protein.

Given these unexpected characteristics of PrgE, it is tempting to speculate about its evolutionary origin. Despite being present in the middle of a T4SS operon on a bacterial conjugative plasmid, PrgE does not behave at all like a bacterial SSB. No close structural homologs could be identified via Dali^46^ and Foldseek^45^. PrgE’s oligomerization behavior in DNA-binding, where PrgE monomers can be added like beads on a string in a noncooperative manner, is reminiscent of some viruses whose SSBs have a monomer as a functional subunit that can be added on ssDNA^26,55^. We did find similarities regarding DNA-binding affinities with an archaeal SSB, which is described as resembling viral SSB-like proteins^54,56^. Indeed, the C-terminally truncated Enc34 phage SSB has been shown to bind dsDNA^57^. Furthermore, the Enc34 SSB was also suggested to be able to bind DNA in a filamentous manner, similar to what we here observe for PrgE^57^. Additionally, PrgE was originally annotated as an SSB protein based on its 37 % sequence similarity to a lactococcal phage SSB^16^. We therefore find it likely that PrgE at some point has been introduced to pCF10 via horizontal gene transfer mediated by a phage.

What then is the function of PrgE for the T4SS and in conjugation? PrgE is expressed as part of the P_Q_ operon of pCF10, surrounded by proteins that are essential for its T4SS (**Fig. 1**). This means that PrgE will be produced only when transcription of the P_Q_ operon has been induced, and its production will be quickly shut down again, just like the rest of the proteins encoded by the P_Q_ operon^12^. Our first hypothesis was that PrgE might interact with other important DNA-binding components of type IV secretion, the relaxosome proteins PcfG and PcfF, as SSBs can be important players for recruiting proteins to DNA^58,59^. However, PrgE does not seem to interact strongly with either of them. Secondly, we speculated that PrgE was important for conjugation in other ways, potentially by protecting the conjugative ssDNA in either the donor or recipient strain, or maybe by aiding the establishment of the plasmid in the recipient cells^34^. To test this, we created a knock-out of PrgE (pCF10:Δ*prgE*). However, no significant differences in conjugation efficiency could be observed, neither in exponential phase nor in stationary phase. It also did not affect the efficiency during multiple serial conjugation events. This is in line with what was observed in previous studies on a F-plasmid, where knocking out a plasmid encoded *ssb* also did not reduce mating rates^35^. However, these experiments were performed under lab conditions, and it is possible that PrgE does contribute to conjugation efficiency under other, less ideal, circumstances.

Conjugative plasmids retain many proteins that are not strictly required for conjugation itself, but provide various other advantages, for example competitiveness against other conjugative elements or replacement of host functions that allows plasmids to use a wider host range^15^. One potential avenue is to explore if PrgE suppresses the SOS response in recipient cells like the F-plasmid SSB does^35^. However, we deem it unlikely that PrgE has a homologous function, given that F-plasmid SSB is a typical bacterial SSB that can compensate for genomic SSB deficiencies^60,61^, while PrgE is very different from *E. faecalis* SSB and has very unusual DNA-binding characteristics. Understanding the exact function of PrgE remains an exciting prospect for future research.

Conjugative plasmids have been studied for many decades now, ever since the R1 conjugative plasmid was first isolated from a clinical isolate in 1963^62^. Genes encoding for OB-fold proteins are part of these plasmids, but our understanding of their specific function within conjugation remains very limited and is almost exclusively based on T4SSs from Gram-negative bacteria. Here, we have shown that PrgE from the Gram-positive conjugative plasmid pCF10 behaves differently to the more well-studied SSBs. It binds ssDNA by attaching PrgE monomers to the DNA like beads on a string, instead of around a globular oligomer like E. coli SSB, and it binds dsDNA equally well as ssDNA. Its oligomerization behavior and DNA-binding mechanism is instead providing insight into a class of OB-fold proteins that has been very poorly characterized.

## Acknowledgments

The authors would like to thank Dr. Krishna Chaitanya Bhattiprolu, Dr. Lena Lassinantti and Dr. Saba Shahzad for input on PcfG production and purification. We also thank Dr. Josy ter Beek for rewarding discussions regarding the project. We acknowledge Protein Production Sweden (PPS) for providing facilities and experimental support. PPS is funded by the Swedish Research Council as a national research infrastructure. We acknowledge MAX IV Laboratory for time on Beamline BioMax under Proposal 20180236. Research conducted at MAX IV, a Swedish national user facility, is supported by the Swedish Research council under contract 2018-07152, the Swedish Governmental Agency for Innovation Systems under contract 2018-04969, and Formas under contract 2019-02496. We also acknowledge the synchrotron ESRF (France) for time at beamlines ID23 and ID30. This work was supported by grants from the Swedish Research Council (2016-03599 & 2023-02423 to R.P-A.B, 2019-01874 to P.H.W.), Knut and Alice Wallenberg Foundation (to each R.P-A.B and P.H.W.) and Kempestiftelserna (SMK-1762 & SMK-1869 to R.P-A.B.)

## Supplemental Figures

**Figure S1.**
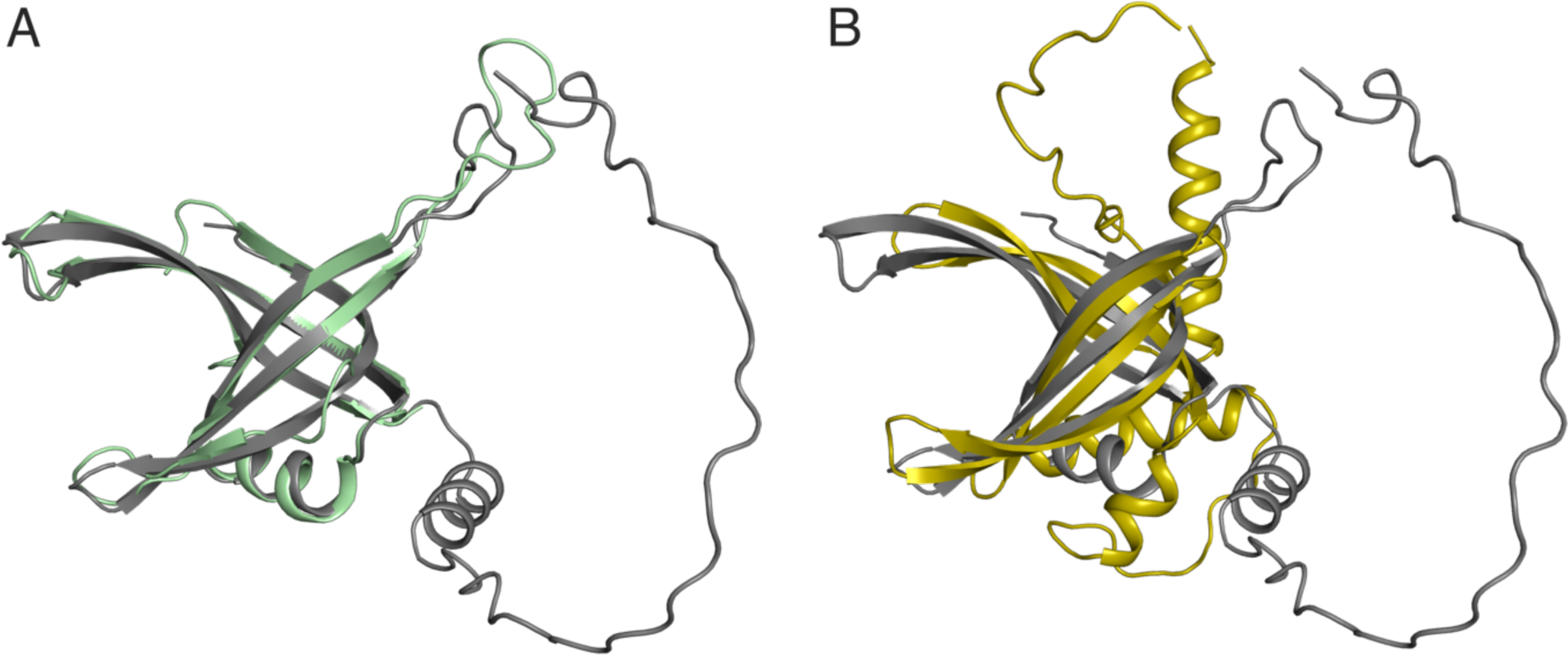
AlphaFold2 model of PrgE differs from bacterial SSBs. **A)** Superimposition of an AlphaFold2 model of genome-encoded *E. faecalis* SSB (accession code: WP_002393727, grey) with *E. coli* SSB (PDB: 1EYG, green). **B)** Superimposition of AlphaFold2 models of PrgE (yellow) and genome-encoded *E. faecalis* SSB (grey).

**Figure S2.**
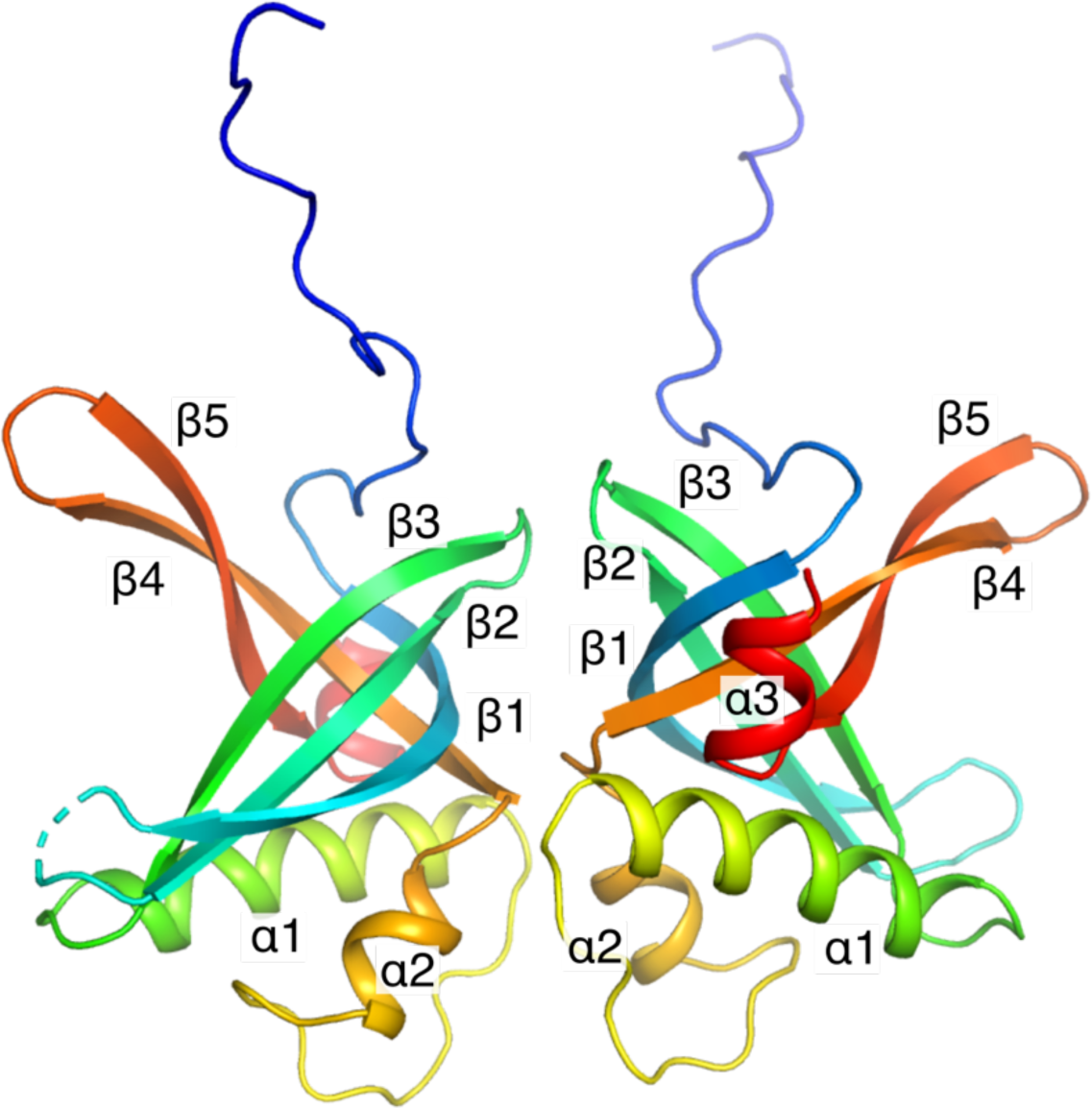
Structure of apo PrgE. Two PrgE molecules are found in the asymmetric unit of the crystal, potentially showing a dimeric form of PrgE when it is not bound to any nucleic acid.

**Figure S3.**
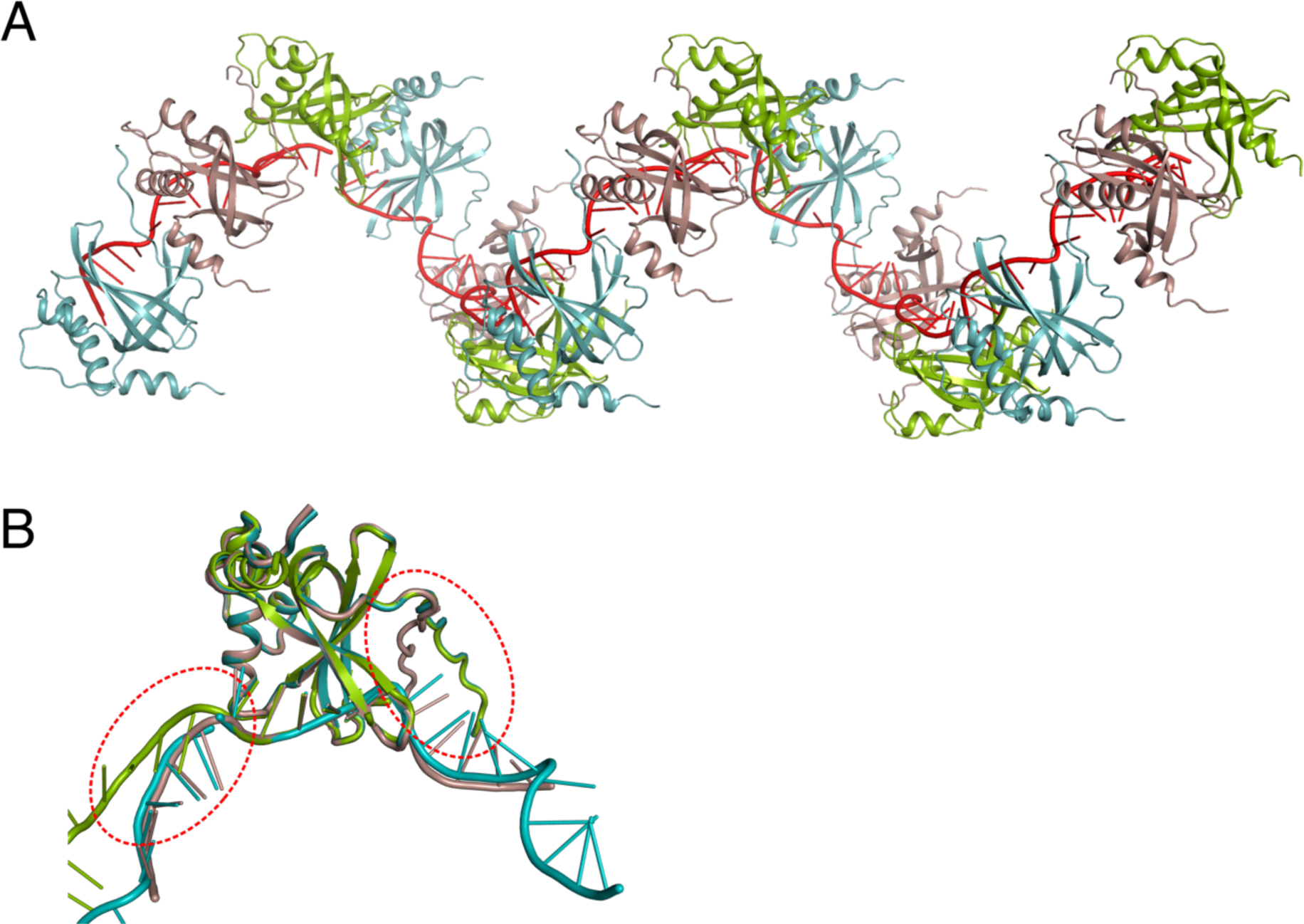
ssDNA forms a continuous strand throughout the crystal packing. **A)** Structure of 5 asymmetric units (each containing 3 PrgE molecules, subunit A in teal, subunit B in brown and subunit C in green) bound to a total of 75 DNA bases highlighted in red. **B)** Superimposition of the three different chains of the PrgE structure bound to ssDNA. Red circles highlight the different angle in the DNA kink in subunit C and the corresponding different angle of the N-terminal tail in subunit B.

**Figure S4.**
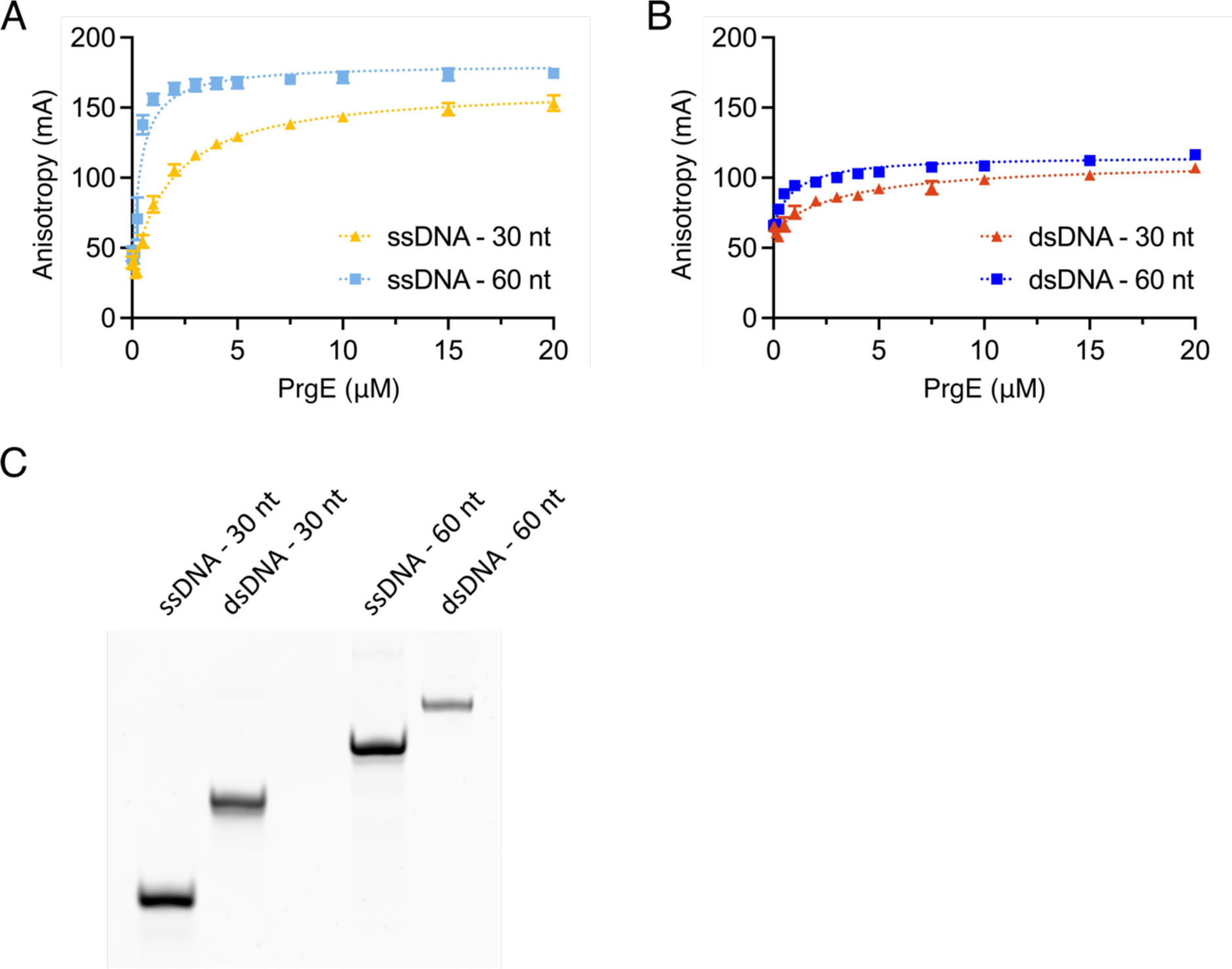
Fluorescence anisotropy measurements of PrgE (0 to 20 *μ*M) in reaction buffer with ssDNA or dsDNA in buffer containing 100 mM NaCl. **A)** PrgE binding to 30-mer (yellow) and 60-mer (light blue) ssDNA substrates. **B)** PrgE binding to 30-mer (orange) and 60-mer (blue) dsDNA substrates. Error bars represent the standard deviation (n=3). **C)** Native gel analysis of the ssDNA and dsDNA substrates used for fluorescence anisotropy.

## Supplemental Tables

**Table S1.**
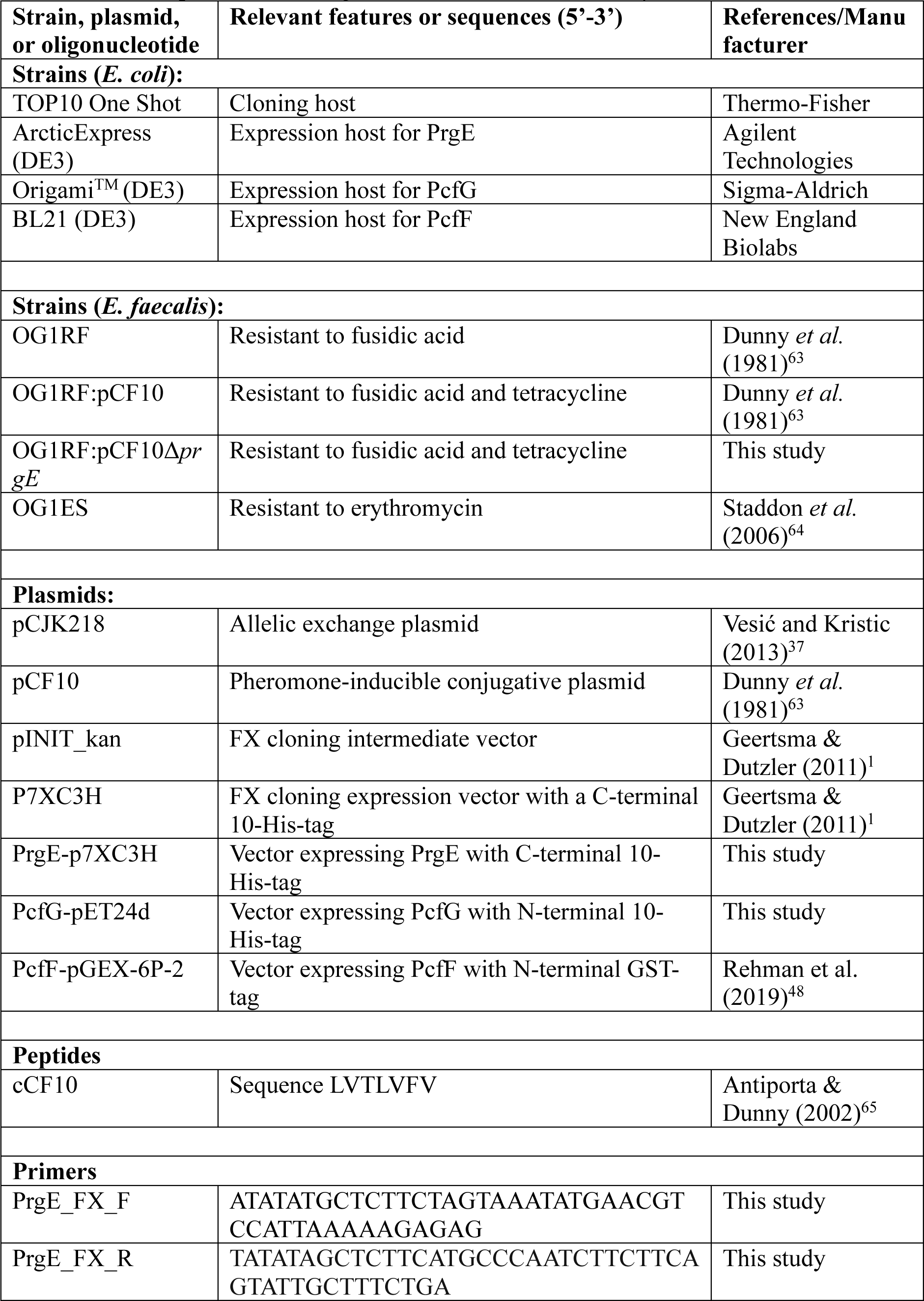
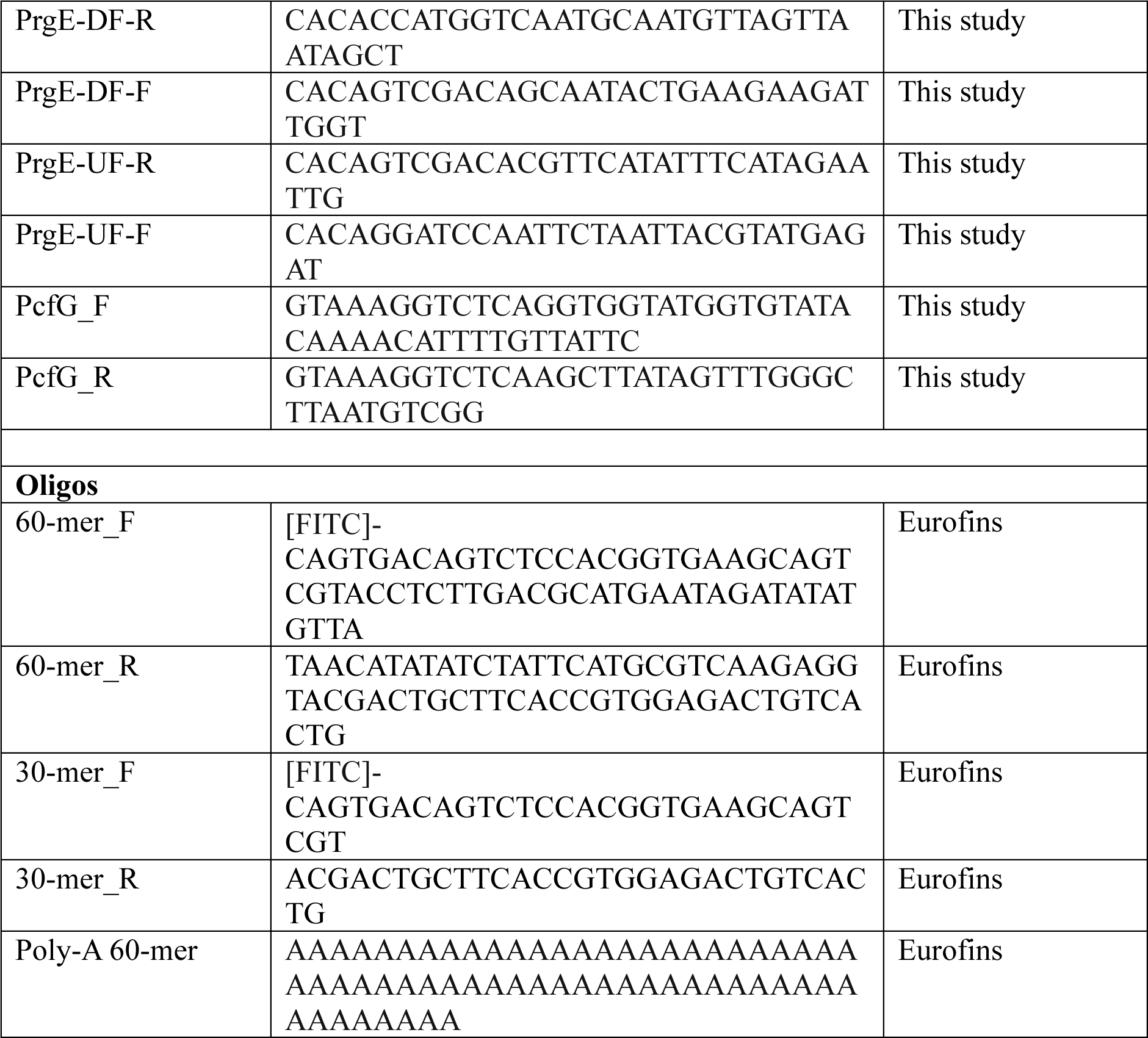
Strains, plasmids and oligonucleotides used in this study.

**Table S2.** Data collection and refinement statistics.

| **Data collection summary** | **PrgE apo structure** | **PrgE DNA-bound** |
| --- | --- | --- |
| Resolution range | 48.38 – 2.51 (2.56-2.51) | 49.74 - 2.67 (2.77 - 2.67) |
| Space group | P 2<sub>1</sub> 2<sub>1</sub> 2<sub>1</sub> | P 2<sub>1</sub> 2<sub>1</sub> 2<sub>1</sub> |
| Cell dimensions |  |  |
| a, b, c (Å) | 58.175 58.17 87.099 | 75.776 90.043 131.848 |
| α, β, γ (°) | 90 90 90 | 90 90 90 |
| Total reflections | 347213 (18816) | 175730 (17998) |
| Unique reflections | 10486 (537) | 26266 (2536) |
| Multiplicity | 33.1 (35.04) | 6.7 (7.0) |
| Completeness (%) | 99.6 (100) | 99.75 (99.06) |
| Mean I/sigma (I) | 21.4 (2.39) | 10.71 (0.95) |
| R-meas | 13.9 (1.718) | 0.1158 (2.067) |
| CC(1/2) | 0.999 (0.852) | 0.998 (0.628) |
| **Refinement summary** |  |  |
| Resolution range | 48.38-2.7 (2.77-2.7) | 49.74 - 2.67 (2.74 - 2.67) |
| R-work | 0.2345 (0.327) | 0.2305 (0.468) |
| R-free | 0.2777 (0.419) | 0.2523 (0.492) |
| Number of non-hydrogen atoms | 2165 | 3716 |
| protein | 2135 | 3372 |
| DNA |  | 315 |
| other ligands | 11 | 20 |
| solvent | 19 | 9 |
| RMS (bonds) | 0.002 | 0.006 |
| RMS (angles) | 0.67 | 1.05 |
| Ramachandran favored (%) | 98.02 | 96.52 |
| Ramachandran allowed (%) | 1.98 | 3.48 |
| Ramachandran outliers (%) | 0.00 | 0.00 |
| Average B-factor | 64.0 | 95.76 |
| protein | 64.29 | 94.37 |
| DNA |  | 111.27 |
| other ligands | 101.09 | 99.49 |
| solvent | 49.62 | 63.43 |

Statistics for the highest-resolution shell are shown in parentheses.

**Table S3.**
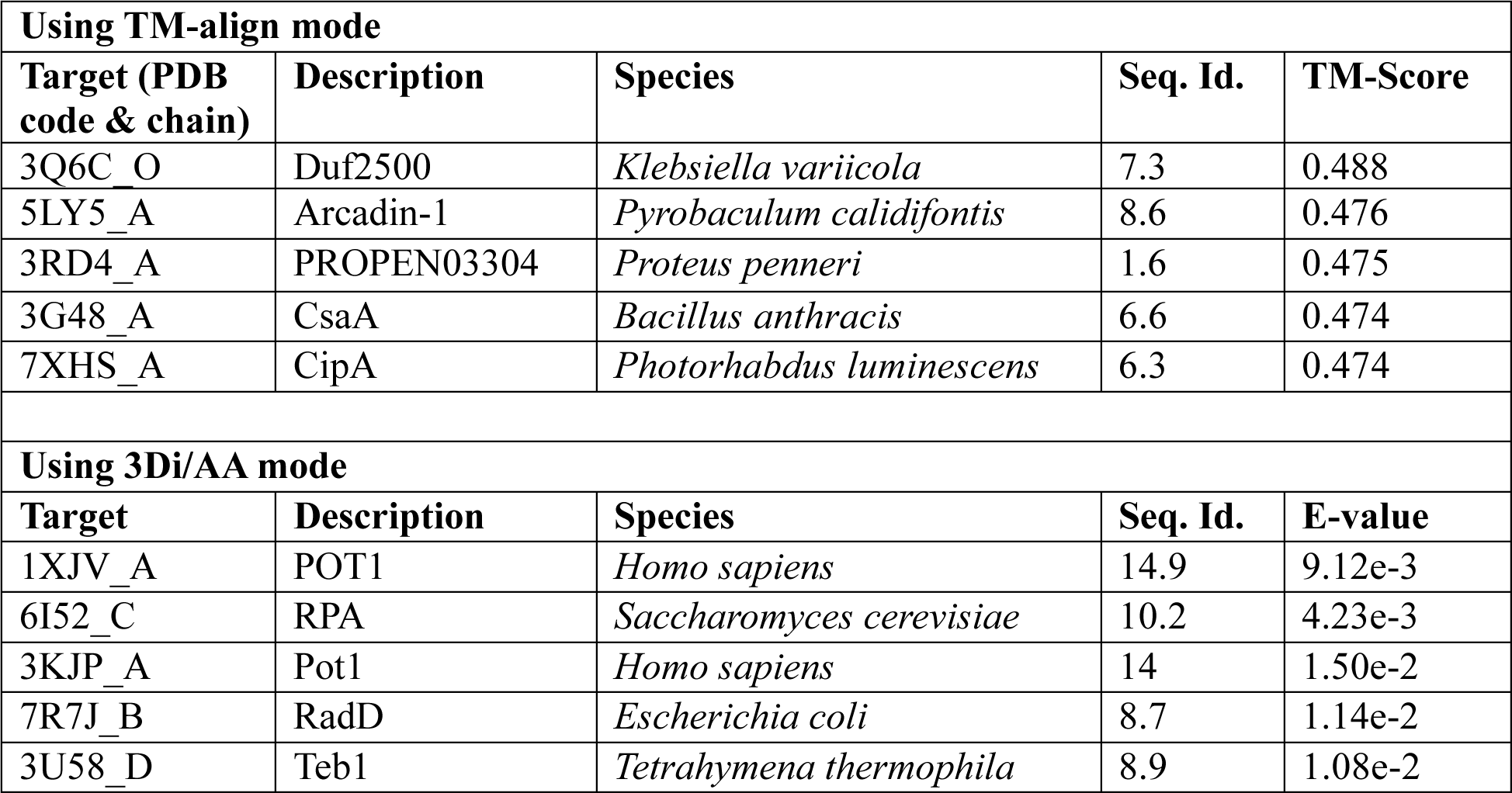
Structural homology searches using Foldseek reveal low homology to other characterized proteins.

## Notes

### Competing Interest Statement

The authors have declared no competing interest.

